# SARS-CoV-2 Variant-Dependent Alterations in Nasopharyngeal Microbiota and Host Inflammatory Response

**DOI:** 10.1101/2025.10.30.685528

**Authors:** Mario Maulen, Ignacio Pezoa-Soto, Daniel León, Ela M. Dominguez, Alberto J.M. Martin, Waldo A. Díaz-Vásquez

## Abstract

The SARS-CoV-2 pandemic saw multiple outbreaks occur over short periods. This was linked to the virus’s high infectivity and rapid mutation rate, which hindered the development of effective treatments and a comprehensive understanding of COVID-19 pathophysiology.

The nasopharyngeal tract, the main entry site of SARS-CoV-2, interacts with angiotensin-converting enzyme 2 (ACE2) receptors via the viral spike glycoprotein, triggering pro-inflammatory responses that affect both tissue integrity and the resident microbiota. Viral infection induces dysbiosis by modulating microbiome diversity. Although distinct variants produce independent symptoms and viral loads, their impact on the nasopharyngeal microbiota and host inflammatory profile remains poorly understood.

To address this, we analyzed nasopharyngeal samples from Chilean individuals collected during different phases of the pandemic. The microbiota was characterized by 16S rRNA gene sequencing (V3–V4 region), and cytokine expression was quantified by RT-qPCR. Finally, applied rigorous data processing and machine-learning models Random Forest and K-Nearest Neighbors to identify associations between SARS-CoV-2 variants, inflammatory markers, and opportunistic bacterial genera.

Our results reveal that SARS-CoV-2 infection promotes distinct immunomicrobial signatures marked by TNF-α-driven inflammation and the expansion of opportunistic taxa. These variant-dependent alterations indicate that host inflammatory responses and microbial dysbiosis are closely intertwined in the nasopharyngeal environment. This study provides a comprehensive framework integrating microbial and host factors to better understand the mechanisms underlying COVID-19 pathogenesis and highlights the potential of combining microbiome and cytokine profiling with machine-learning approaches to differentiate infection outcomes across viral variants.

**Importance:** Understanding how SARS-CoV-2 infection alters the upper respiratory microbiota and host immune responses is essential to uncover the mechanisms underlying COVID-19 severity. In this study, we integrate bacterial 16S rRNA sequencing, inflammatory markers, and machine learning to identify key descriptors of infection such as TNF-α, Acinetobacter, Prevotella, and Staphylococcus. These markers define a conserved immunomicrobial signature across viral variants, linking inflammatory activation with microbial dysbiosis in the nasopharyngeal environment. Our findings highlight the value of the correct and robust application of machine learning models and nested cross-validation strategies in high-dimensional (wide) datasets to improve variant discrimination, reveal host–microbe interactions, and support the development of therapeutic strategies for respiratory viral infections.

## Introduction

During the COVID-19 pandemic, scientific research focused on the infection mechanisms of SARS-CoV-2 as a potential therapeutic strategy. The virus’s high infectivity, combined with a rapid mutation rate, led to the emergence of multiple variants of concern and interest. These variants have been grouped into clades and lineages based on the accumulation of conserved point mutations and deletions over time^1^. To handle the increasing cases of hospitalized subjects and the collapse of the healthcare system, mRNA-based vaccines targeting SARS-CoV-2 structural proteins were developed, each exhibiting varying degrees of effectiveness. Most of these vaccines focus on the spike protein, a viral membrane glycoprotein composed of two subunits (S1 and S2), which mediate viral attachment and membrane fusion with host cells expressing the ACE2 receptor^2^. The infection process begins when the S1 subunit binds to ACE2 via its receptor-binding domain, located on the outermost surface of the protein. This domain enables the virus to anchor to host cells with high affinity^3,4^. A subsequent cleavage between the S1 and S2 subunits is mediated by the serine transmembrane protease TMPRSS2 at a multibasic cleavage site on the spike protein^5^. This triggers a conformational change in the S2 subunit, allowing it to elongate and insert into the host cell membrane, initiating membrane fusion and infection^6^. Once inside the host cell, SARS-CoV-2 releases its RNA genome into the cytoplasm, where a viral RNA-dependent RNA polymerase replicates the genome for translation, glycosylation, assembly, and eventual release of new virions^7^.

The respiratory tract remains the primary target of SARS-CoV-2, where viral titers are typically highest, and severe symptoms can lead to fatal outcomes^8^. Recent evidence indicates that SARS-CoV-2 infection also modulates the respiratory microbiota; however, most studies to date have focused on only one variant, primarily the earlier variants of concern^9^.

Given that the nasopharynx is the first region of the human respiratory system exposed to the virus, opportunistic and co-infecting pathogens have been identified at this site in both mild cases and patients requiring mechanical ventilation^10^.

Differences in nasopharyngeal microbiota composition between healthy individuals and COVID-19 patients have been documented in various studies, revealing reductions in commensal bacterial populations and increases in opportunistic pathogens^11,12,13^. Similar alterations have been reported for the influenza virus, where the respiratory microbiome is reshaped at multiple anatomical sites during both the acute and recovery phases of infection^14^. Such dysbiosis, or microbial imbalance, resulting from SARS-CoV-2 infection, directly affects host tissues due to elevated levels of pro-inflammatory factors such as toxins and pathogen-associated molecular patterns^15^. Studies of the upper respiratory tract in COVID-19 patients have shown elevated local expression of TNF-α and IFN-γ^16^. Interleukin-2, a key cytokine in CD8+ T-cell activation, has been positively correlated with antibody titers in infected individuals^17^; however, early-phase overexpression can trigger a cytokine storm, significantly worsening the clinical outcome^18^. Additionally, increased IL-1ra levels have been reported in critically ill subjects, possibly indicating a compensatory anti-inflammatory response to excessive immune activation driven by opportunistic bacteria^19^. The multivariable nature of this phenomenon complicates efforts to define which variables are dependent or independent in the pathway from viral infection to microbial dysbiosis and the associated inflammatory response.

These challenges emphasize the need for computational tools capable of identifying often non-obvious associations among heterogeneous variables, particularly in datasets constrained by small sample sizes^20^. In this context, machine learning (ML), has emerged as a valuable approach for identifying, classifying, and analyzing complex patterns that would otherwise remain undetected^21^. ML has been widely applied in microbiome and sequencing-based research^22,23,24^. Among supervised learning models, Random Forest (RF) and K-Nearest Neighbors (KNN) are prominent due to their ability to handle non-linear relationships and infer associations from high-dimensional data by using distance-based methods like Euclidean distance in KNN^25, 26^. These models outperform traditional linear approaches when it comes to complex biological systems.

However, a major limitation of such algorithms is the availability of sufficient data to ensure the reliability of the inferred relationships^27^. Data shortage may arise from limited sample collection, insufficient resources to generate omics data per sample, or difficulties in recruiting subjects with specific characteristics. This challenge is further compounded by the high dimensionality of omics datasets, which creates additional hurdles for applying ML. These so-called “wide datasets” require specific protocols to enhance the robustness of inferred associations^28^. Despite these difficulties, and some ongoing debates, ML approaches applied to “wide datasets” have shown great potential for generalization across various domains, such as identifying molecular properties and expression patterns in distinct tumor types^29^. ^30, 31, 32^.

Considering that the symptomatology associated with COVID-19 has varied depending on the infecting variant, and that nasopharyngeal dysbiosis and opportunistic pathogen overgrowth alter the host’s inflammatory response, we propose to establish associations between SARS-CoV-2 variants and specific dysbiosis profiles, presence of opportunistic pathogens, and characteristic levels of inflammatory markers.

## Materials and methods

### Data Recollection

#### Study design and sample collection

We collected nasopharyngeal samples between April 2020 and August 2021 to detect SARS-CoV-2 infection and the viral variants. These samples were obtained from four healthcare establishments in the Metropolitan Region of Chile and stored at −80 °C until viral, bacterial, and human genetic material could be extracted. This study was approved by the Scientific Ethics Committee of Universidad San Sebastián, which granted a waiver of patient informed consent.

#### Total RNA/DNA extraction

Between April 2021 and April 2022, SARS-CoV-2 detection assays were conducted using the Mag-Bind® Viral DNA/RNA 96 Kit. This allowed for the identification of positive and negative cases, hereafter referred to as infected and healthy subjects, respectively, yielding a total of 242 positive and 63 negative samples.

#### Variant Identification

Identifying SARS-CoV-2 variants in our samples was crucial for establishing the corresponding classes for our machine learning models. First, detection assays using TaqMan 2019-nCoV Assay/Control were performed to discriminate positive from negative samples. Subsequently, an RT-qPCR genotyping assay using Hydrolysis Probe qPCR Assay was implemented to detect specific spike mutations (P681R, H655Y, and T859N) in a QuantStudio 5 thermocycler. These mutations were used to identify the Delta, Gamma, and Lambda variants, which represented the predominant circulating lineages in Chile during the study period.

#### V3-V4 Regions of 16S rRNA Gene sequencing

The sequencing of V3-V4 hypervariable regions of the bacterial 16S rRNA gene was conducted by the Environmental Sample Preparation and Sequencing Facility (ESPSF) at Argonne National Laboratory, Chicago, IL, USA, using Illumina MiSeq technology and sequencing-by-synthesis chemistry. The primers 515F (5′-GTGCCAGCMGCCGCGGTAA-3′) and 806R (5′-GGACTACHVGGGTWTCTAAT-3′) were employed to amplify 250 bp amplicons. Demultiplexing of the raw sequences was performed using MetaScope (v4.1)^33^. Subsequently, the DADA2^34^ pipeline was applied for quality filtering, dereplication, denoising, and chimera removal, ultimately yielding a phyloseq object for downstream microbial community analysis.

### Microbial Community Analysis in Nasopharyngeal Samples

The richness and abundance of bacterial genera present in each nasopharyngeal sample were determined. To assess statistically significant changes in abundance, DESeq2 package (R v4.1.2) ^35^ was used. Taxonomic assignment of the amplicon sequence variants (ASVs) was performed using the SILVA-138 database^36^. Alpha diversity metrics (Observed, Chao1, Chao2, Simpson Index and Shannon index) were calculated using the microbiome R package^37^ to evaluate the loss of commensal bacteria and the enrichment of opportunistic pathogens.

### Quantification of IL-2, TNFα, IFNγ, and IL-1ra in Nasopharyngeal Samples

To characterize the inflammatory response associated with SARS-CoV-2 infection, the expression levels of IL-2, TNFα, IFNγ, and IL-1ra were quantified by RT-qPCR assays on each sample using specific primers and SYBR Green as the intercalating agent for Cycle Threshold (CT) detection. Gene expression was normalized using GAPDH as housekeeping gene, to calculate the relative expression of each marker. The CT values of the inflammatory markers were normalized to GAPDH to compute the relative expression for each cytokine on a per-sample basis, thereby enabling the generation of individualized inflammatory profiles.

### Association Analysis

Following taxonomic profiling and inflammatory marker quantifications, a matrix was constructed. This matrix comprised the relative abundance of each bacterial genus, the quantified cytokine expression levels, and the corresponding SARS-CoV-2 variant infecting each individual. Based on the sample size, we employed an RF^38^ as classification algorithm focusing on feature importances rather than classification accuracy. The SARS-CoV-2 variant served as the target variable. ASV abundance was encoded (0 =Absent, 50>= Low, 500>=Medium, 1500>=High and 1500< Very High), and cytokine levels were included as normalized continuous numerical features.

#### RF-based Feature Ranking and KNN Testing

We applied a RF classifier to predict viral variants from cytokine and bacterial abundance profiles. Prior to training, features with zero variance were removed and categorical predictors were one-hot encoded. Training was carried out using the scikit-learn implementation of RandomForestClassifier with 10,000 trees using nested cross validation (CV) with 10 K-fold as inner validation and Leave-One-Out cross validation (LOOCV) as outer. Performance was quantified using accuracy, balanced accuracy, micro-averaged precision and recall, in addition to weighted one-vs-one receiver operating characteristic-area under the curve (ROC-AUC). Following training, feature importance was obtained across all trees in each fold. Per-fold importance values were recorded and subsequently aggregated to produce a consensus ranking of predictors. The Top 10 and 20 ranked features obtained with RF were validated using a KNN algorithm trained and evaluated under the same nested CV scheme as the RF^39^.

All analyses were implemented in Python using the scikit-learn library^40^.Given the relatively low computational cost and the exploratory nature of the analysis, an extensive parameter sweep was performed whenever feasible to maximize the robustness of the associations identified.

## Results

### Variant Identification

Once we have finished conducting SARS-CoV-2 viral detection assays on collected nasopharyngeal samples, we identify variants by Hydrolysis Probe qPCR Assay in each remaining positive samples as summarized in Table 1.

**Table 1.**
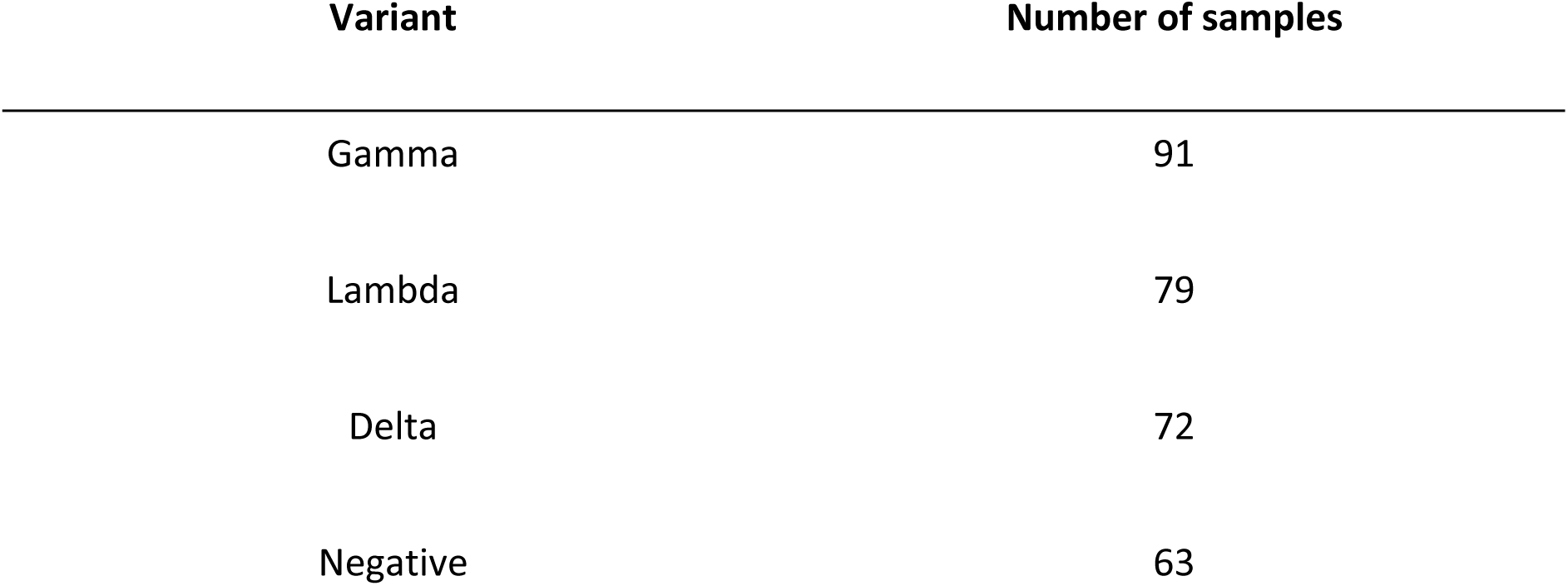
Number of identified samples per variant.

| Variant | Number of samples |
| --- | --- |
| Gamma | 91 |
| Lambda | 79 |
| Delta | 72 |
| Negative | 63 |

### Nasopharyngeal microbiota characterization in Chilean cohort

Analysis of alpha diversity indices revealed significantly lower richness and evenness in SARS-CoV-2–positive samples compared to negatives (Kruskal–Wallis, *p* < 0.05) (Fig. 1), consistent with nasopharyngeal dysbiosis. Within positives, Lambda and Gamma maintained this reduction, while Delta showed diversity values comparable to the negative group. After quality control preprocessing of the sequenced samples, the dominant phyla were identified (Supp. Table 1).

**Figure 1.**
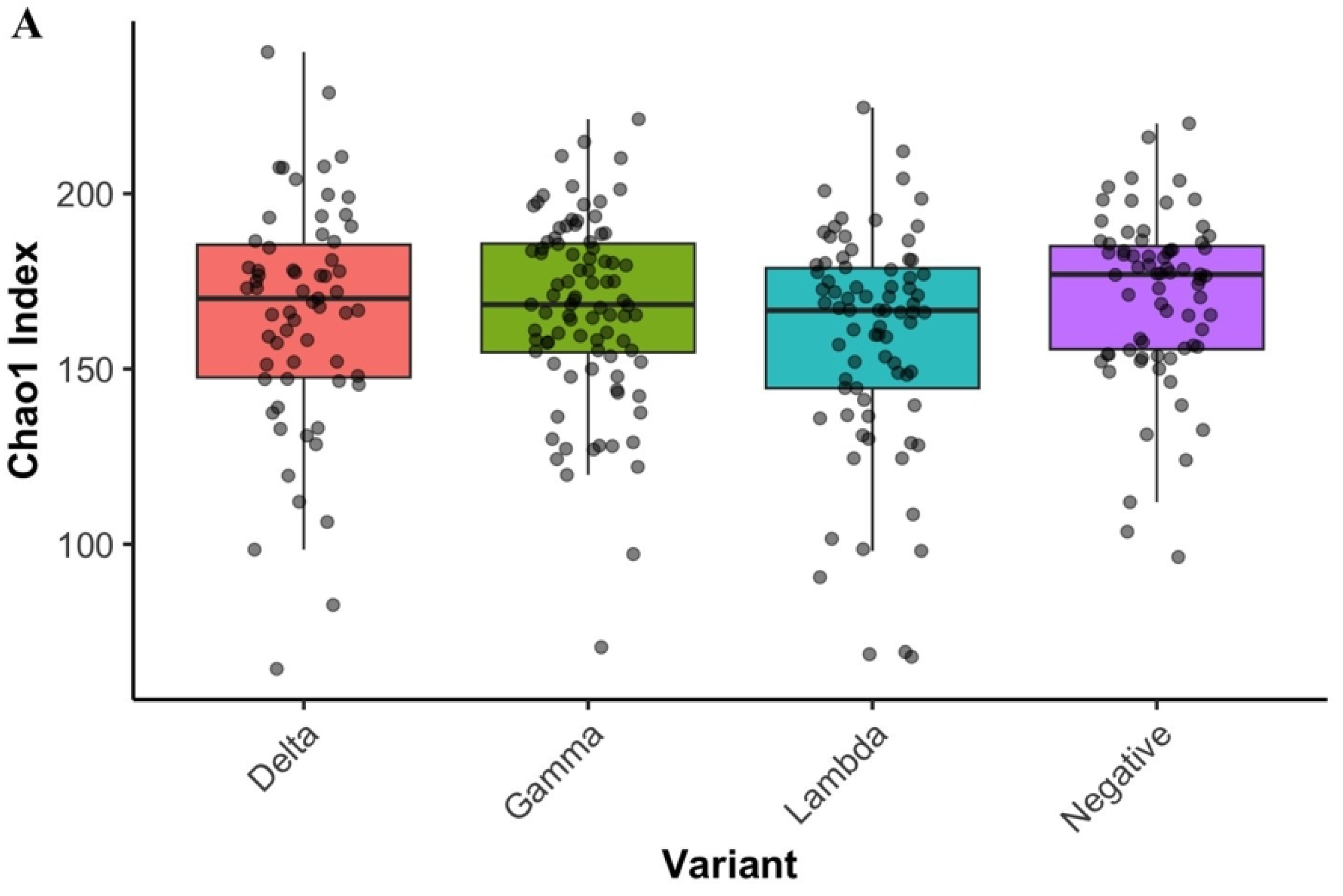
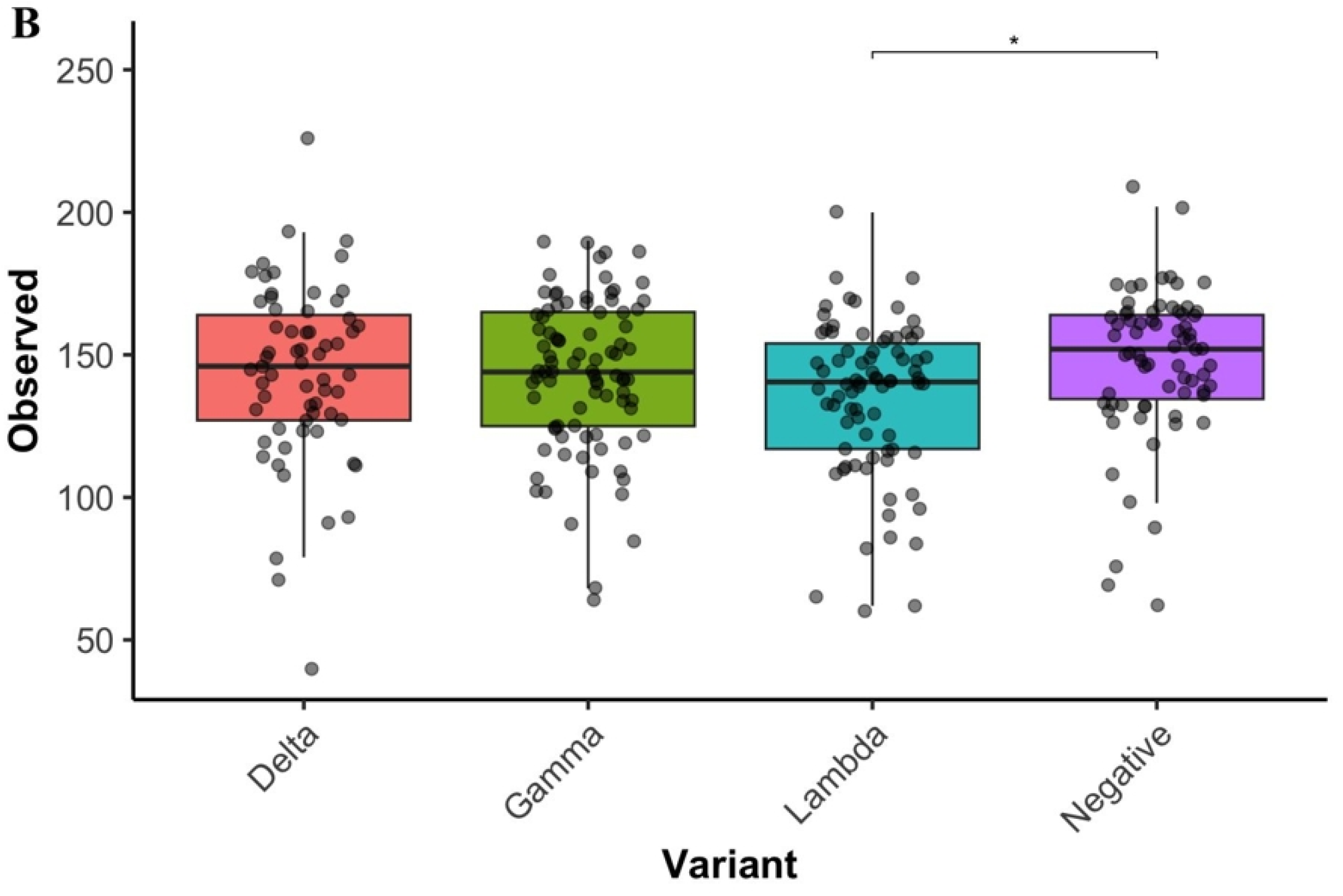
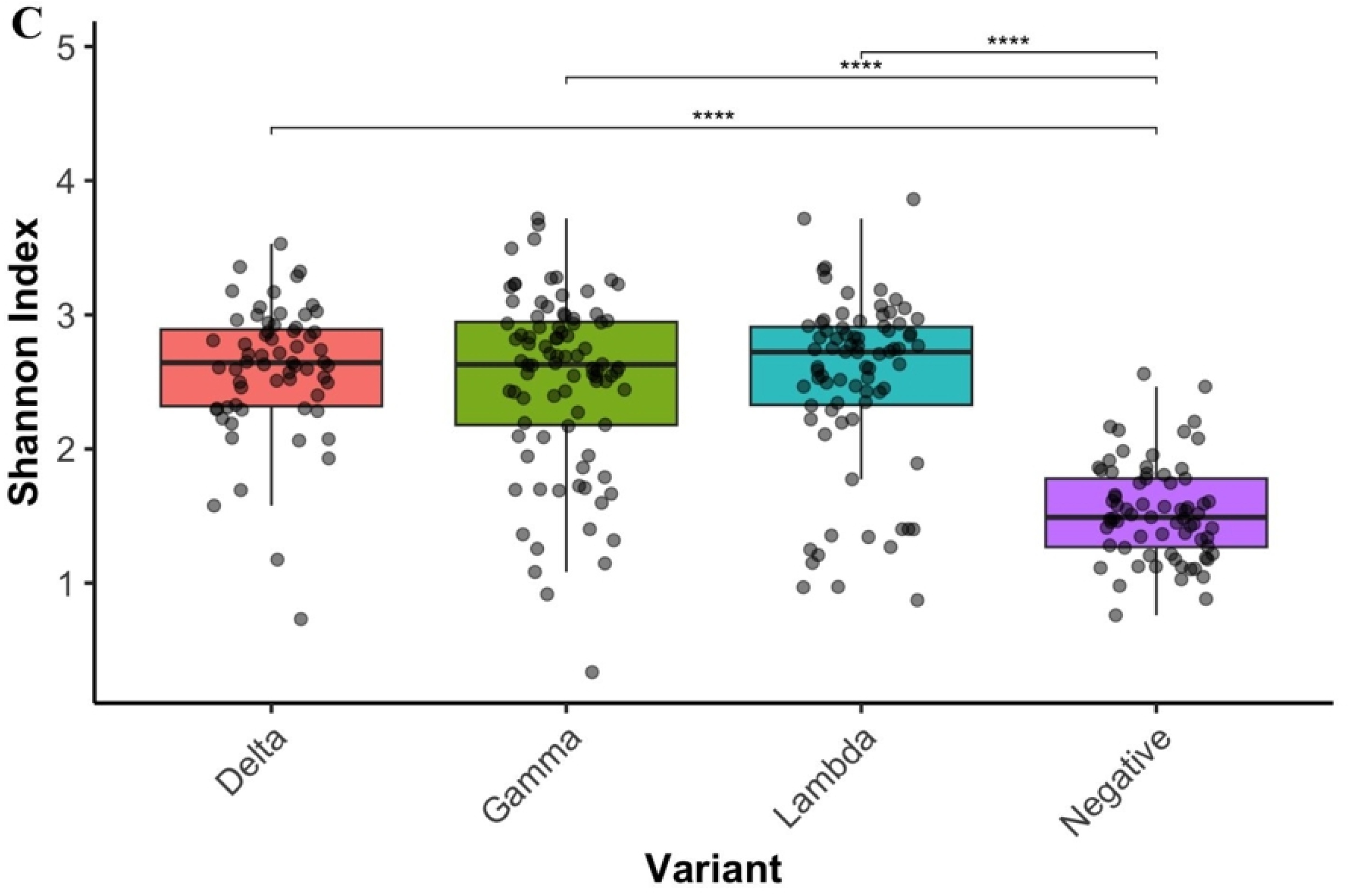
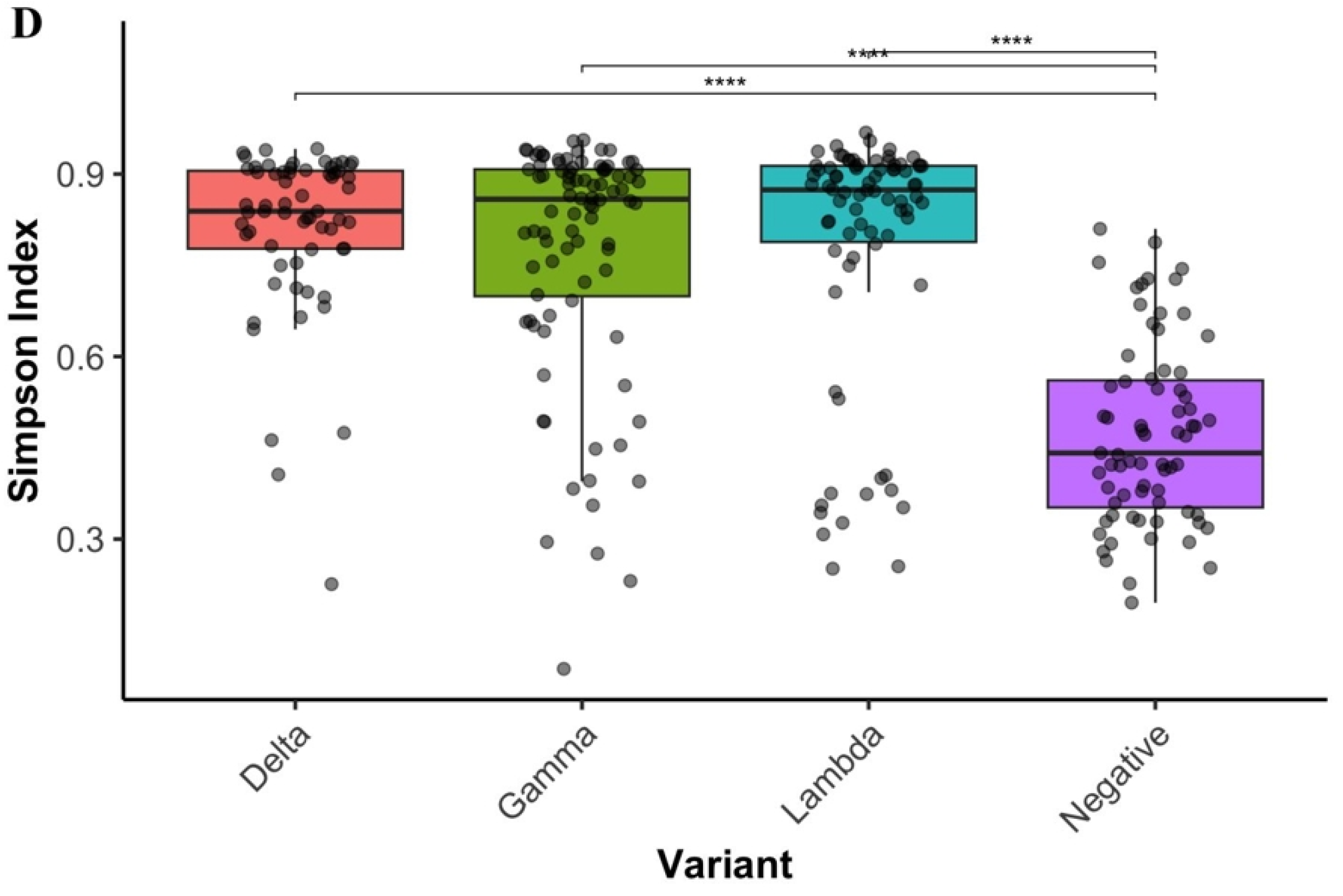
Alpha diversity indices of nasopharyngeal microbiota comparing SARS-CoV-2 positive samples (stratified by Lambda, Gamma, and Delta variants) and negative controls. Box plots display A) of Chao1 richness B), Observed ASVs, C) Shannon diversity and D) Simpson (D) indices. P-values were obtained from Kruskal–Wallis test.

Relative abundance analyses were performed to characterize positive and negative samples to identify the presence and abundance of bacterial genera associated with opportunistic pathogens. We found that the nasopharyngeal microbiota exhibited a modulation between SARS-CoV-2 positive and negative individuals, with compositional and structural shifts that are consistent with microbial dysbiosis in the infected group (Fig. 1). This dysbiotic profile was characterized by altered alpha diversity metrics. Results suggest that SARS-CoV-2 infection disrupts the respiratory microbiome architecture, independent of viral variants. While species richness remains comparable between infected and healthy individuals (Fig. 1A), Shannon and Simpson diversity indices reveal a significant elevation in community evenness among patients infected with Delta, Gamma, or Lambda variants compared to uninfected controls (Fig. 1C-D).

To determine which bacterial taxa abundances were modulated in infected subjects, we analyzed the relative abundance of bacterial phyla and genera. Taxonomic profiling reveals that this disruption manifests as a shift in phylum-level composition, with negative controls exhibiting increased Proteobacteria dominance that is substantially diminished during infection, coinciding with marked expansion of Firmicutes and Bacteroidetes (Fig. 2A). At the genus level, the healthy respiratory microbiome is characterized by clear dominance of Pseudomonas. SARS-CoV-2 infection disrupts this pattern across all variants, triggering the expansion of diverse genera including *Acinetobacter sp., Corynebacterium sp., Dolosigranulum sp., Moraxella sp.* and *Staphylococcus sp.*, thereby transforming the community structure into a more even but dysbiotic state compared to the negative controls (Fig. 2B). This microbiome pattern is similar across all three viral variants examined, suggesting that respiratory dysbiosis represents a conserved feature of SARS-CoV-2 pathogenesis rather than a variant-specific phenomenon.

**Figure 2.**
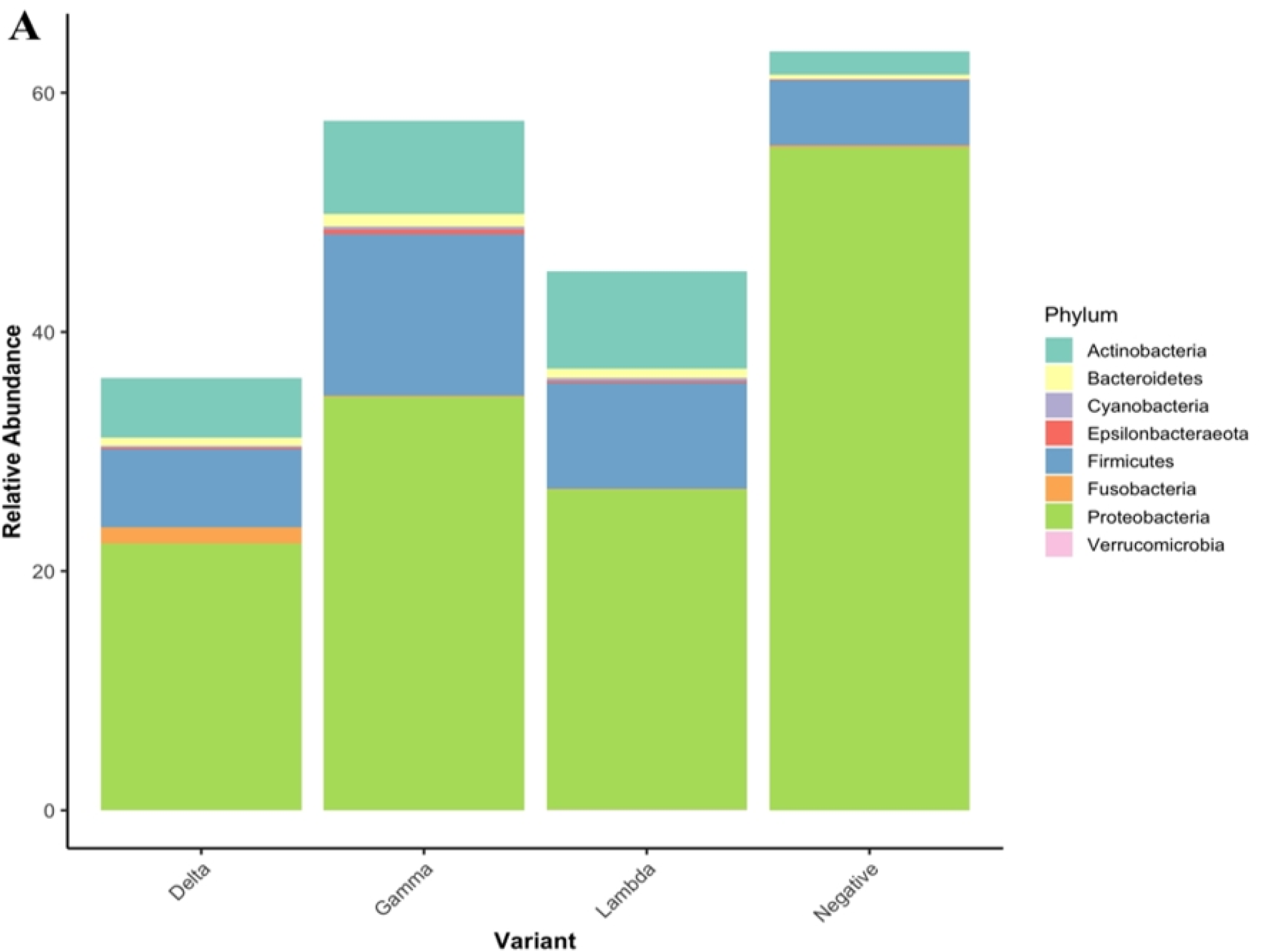
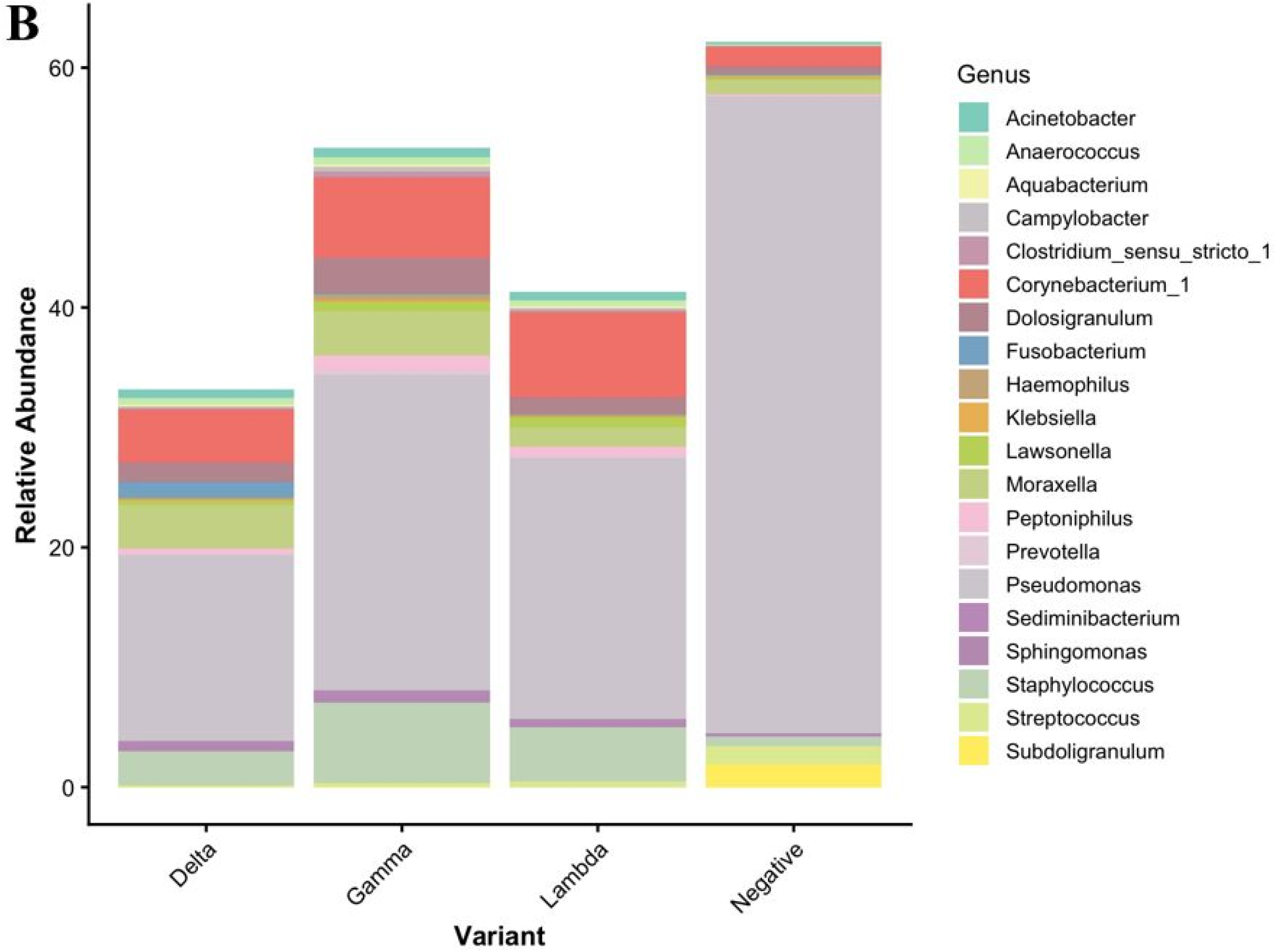
Relative abundance of nasopharyngeal bacterial taxa. Stacked bar plots illustrate the relative abundance of A) bacterial phyla and B) bacterial genera, across SARS-CoV-2 variants and negative subjects.

A more detailed analysis of the most abundant taxa identified a subset of 20 species in the SARS-CoV-2 positive samples, associated with secondary infections in the respiratory tract^41^, such as *Acinetobacter baumannii, Acinetobacter lwoffii*, *Actinomyces odontolyticus, Campylobacter ureolyticus, Corynebacterium jeikeium, Fusobacterium nucleatum, Fusobacterium periodonticum, Haemophilus sputorum, Staphylococcus haemolyticus, Stenotrophomonas maltophilia, Porphyromonas endodontalis, Prevotella buccalis, Prevotella timonensis* and *Prevotella melaninogenica*^13,42,43^.

Once we identified these changes in alpha diversity and the presence of specific phyla and genera in samples from infected individuals, we proceeded to evaluate beta diversity across the different sample groups. Beta diversity analysis revealed significant compositional differences between positive and negative groups, but no distinguishable inter-variant clusters are observed due to the high dispersion and overlap of individual microbial signatures in the ordination space (Fig. 3).

**Figure 3.**
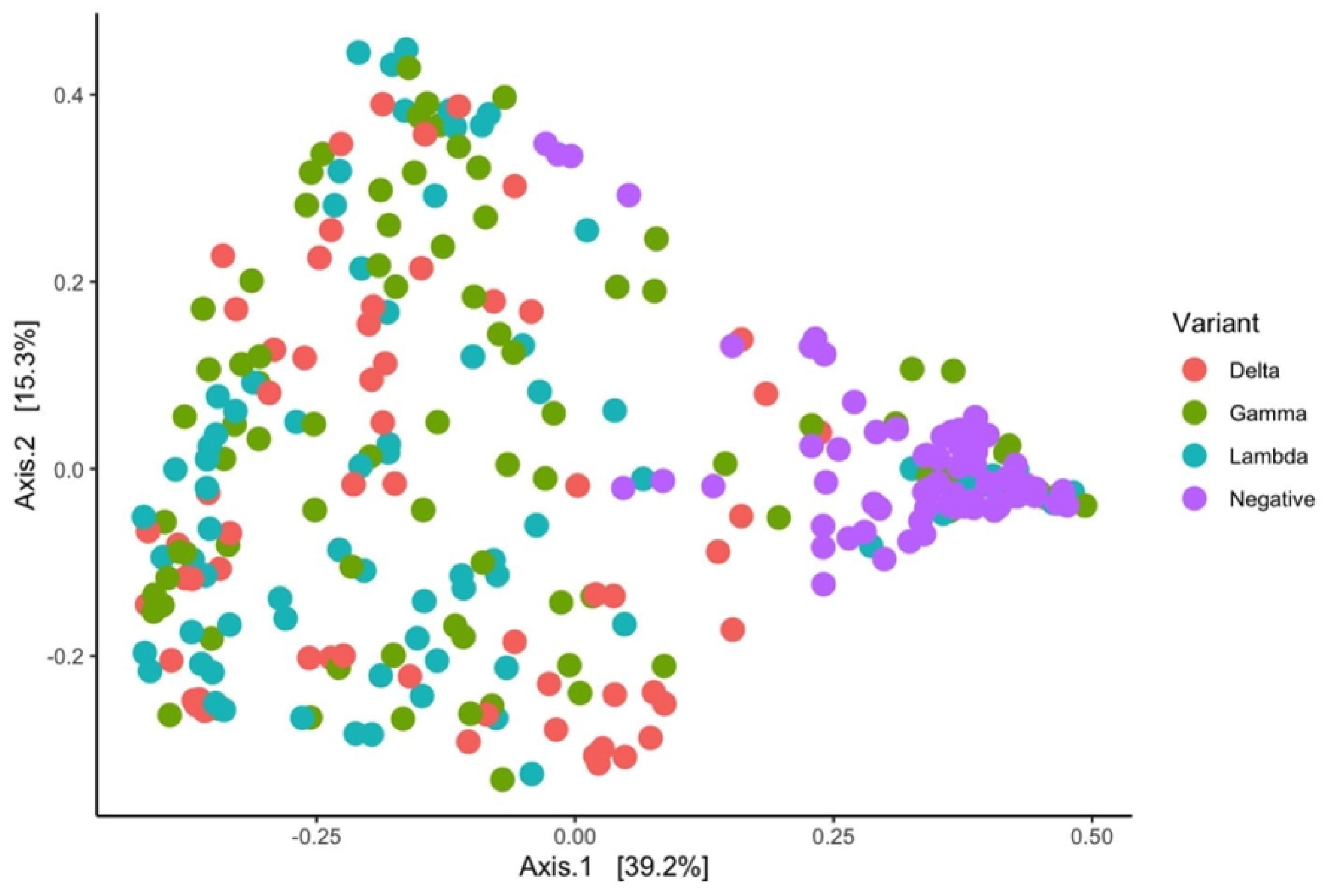
Beta diversity analysis of nasopharyngeal microbiota. Principal coordinate analysis (PCoA) analysis ordination plot based on Bray–Curtis dissimilarity, showing compositional differences between SARS-CoV-2 negative and positive individuals.

Given these observations, we investigated whether such differences or similarities would also be reflected in the inflammatory markers expressed during SARS-CoV-2 infection.

### Inflammatory markers are highly expressed in early variants

The inflammatory profile associated with SARS-CoV-2 infection is highly relevant due to its implications for disease severity, particularly concerning its relation to the cytokine storm^18^, and in asymptomatic patients.

For this reason, expression levels of IL-2, TNF-α, IFN-γ, and IL-1ra were measured to identify potential variations in these inflammatory markers across the dissimilar viral variants included in the study.

Inflammatory marker levels revealed a significant increase in TNF-α expression in SARS-CoV-2–positive subjects compared with negative samples (Fig. 4), consistent with the inflammatory state induced by infection. Other markers showed greater dispersion, suggesting heterogeneous immune responses^44^.

**Figure 4.**
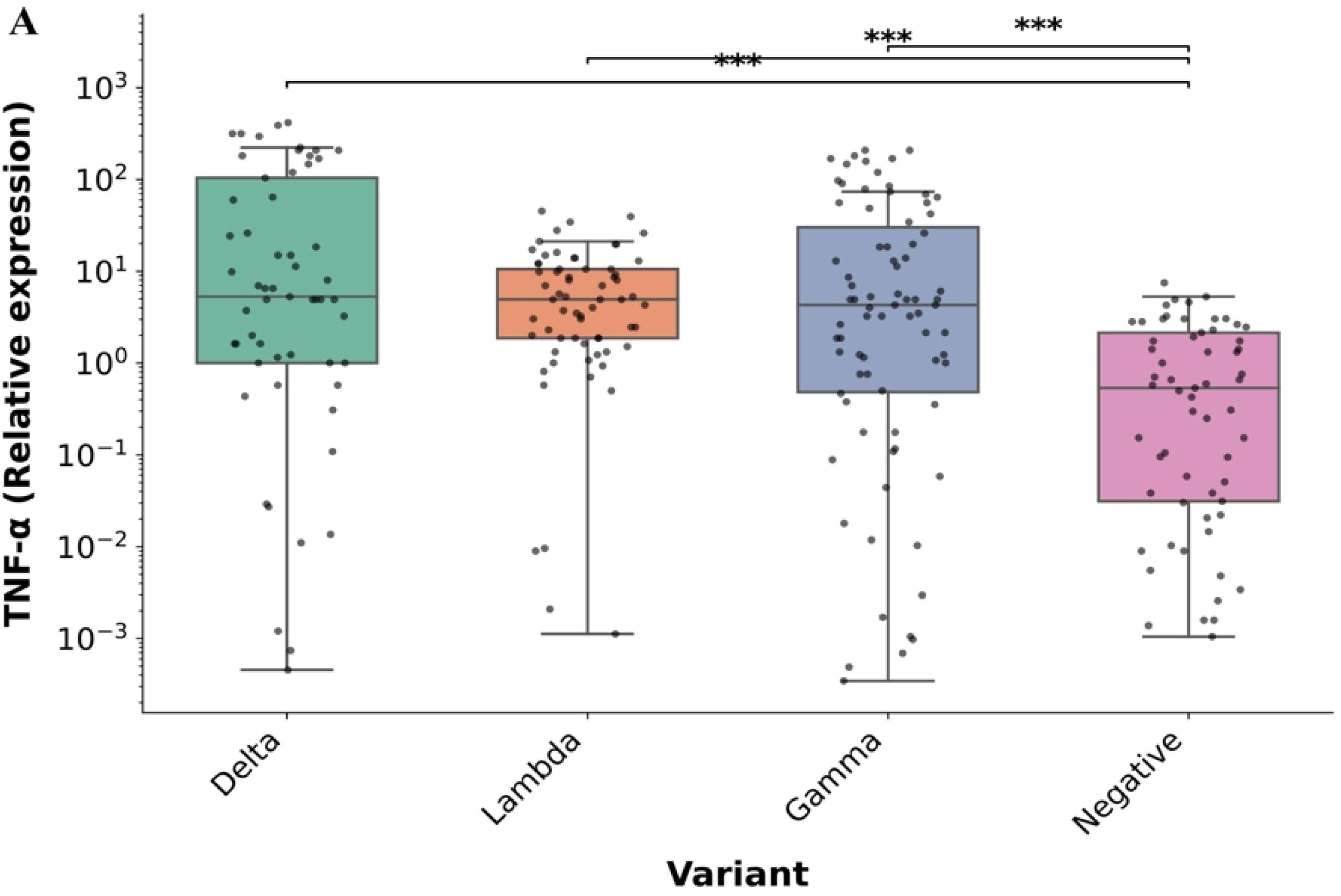
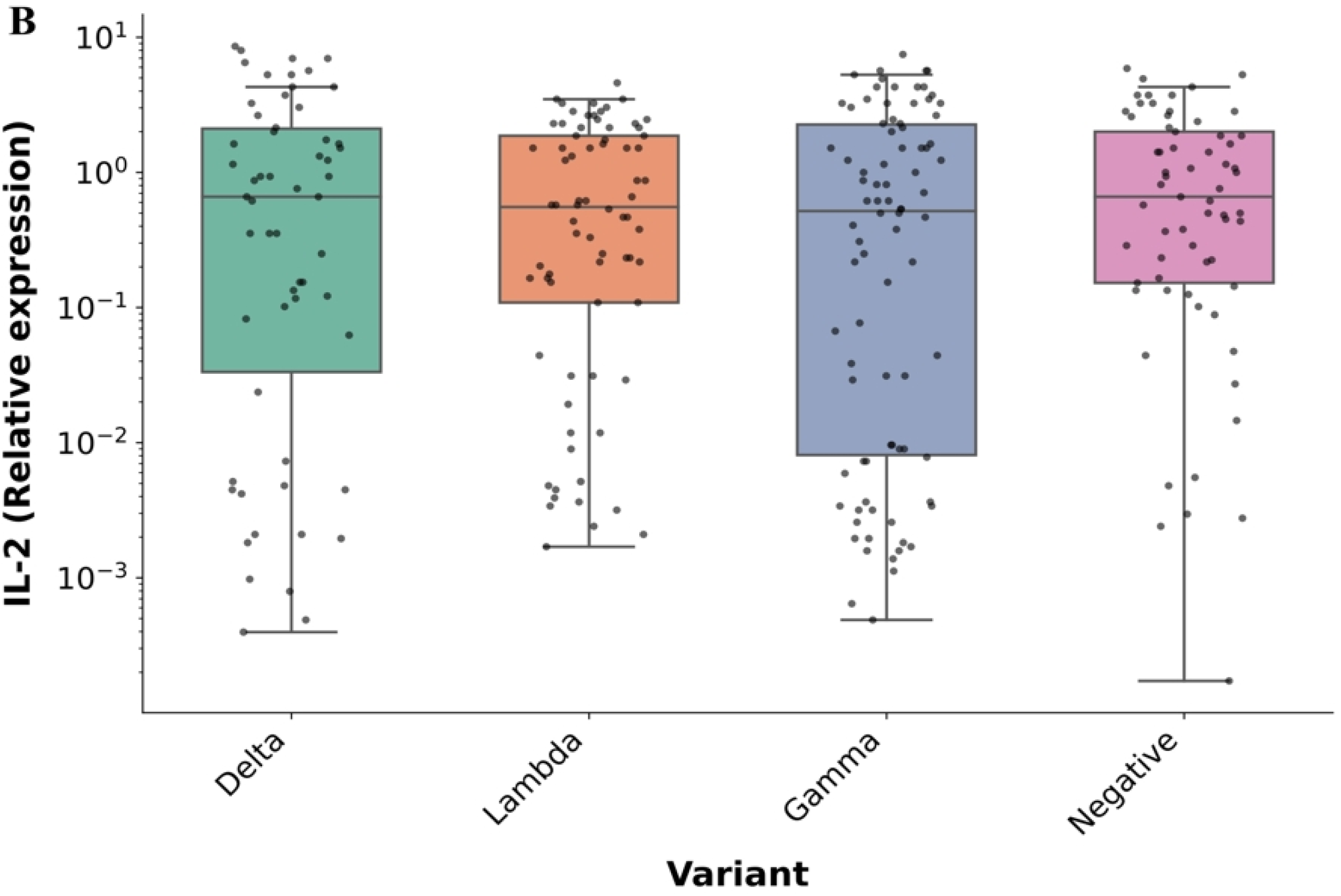
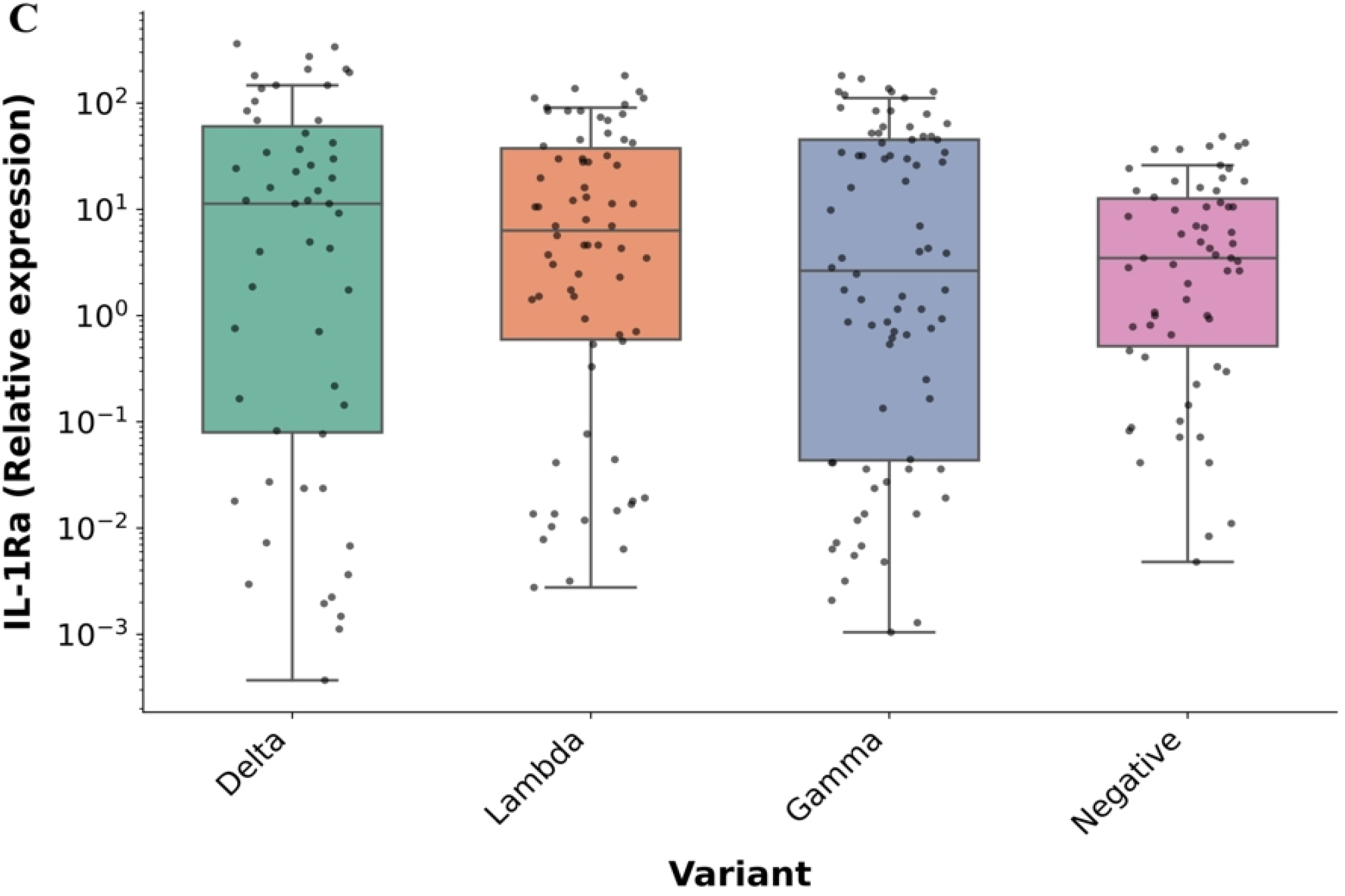
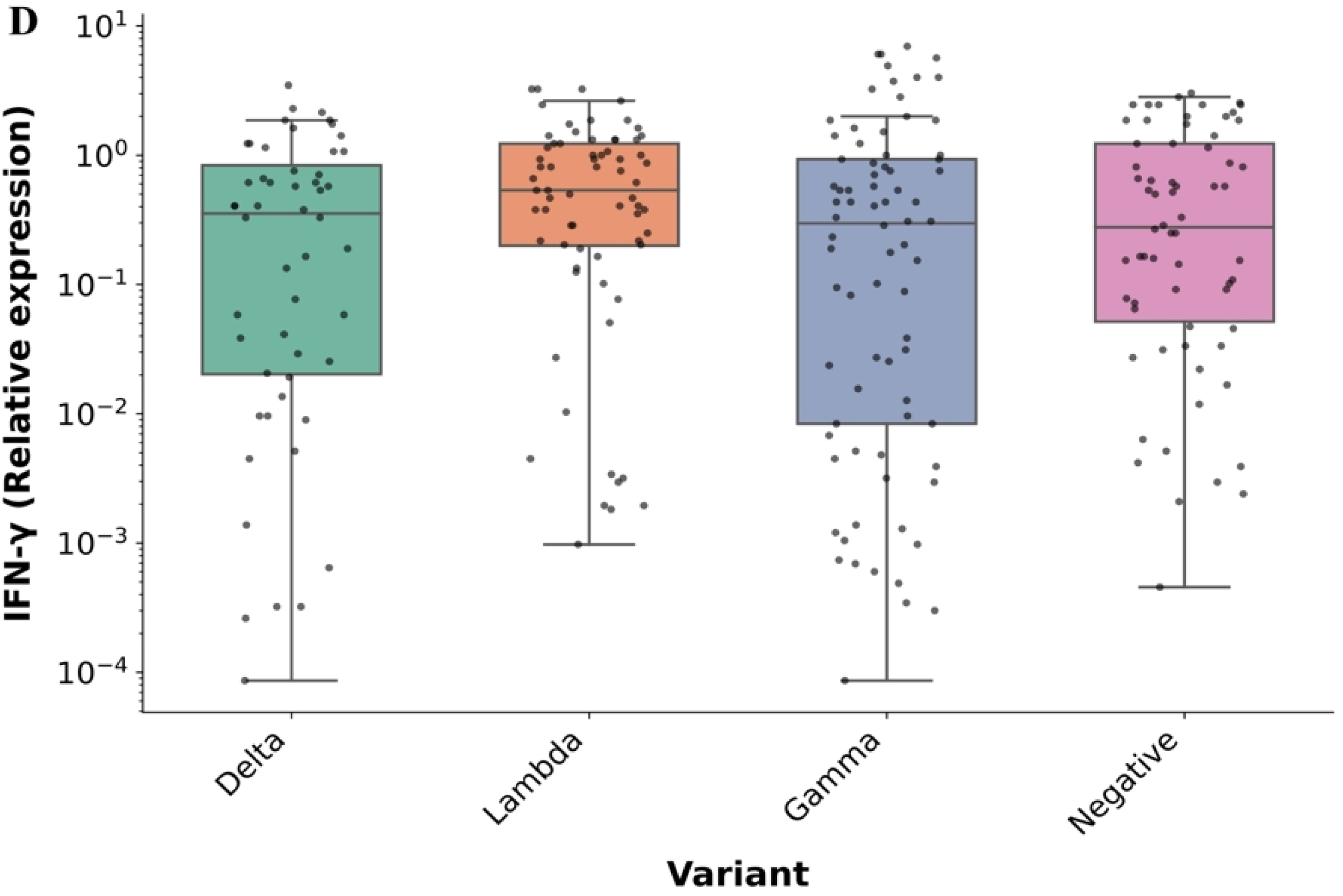
Expression levels of inflammatory markers in SARS-CoV-2 negative and positive subjects. Box plots display the relative expression of A) TNF-α, B) IL-2, C) IL-1ra and (D) IFN-γ normalized to GAPDH. TNF-α expression was significantly higher in positive subjects compared to negatives, with no significant differences observed among viral variants.

### Finding associations based on Machine Learning Models

After completing the experimental assays for nasopharyngeal microbiota analysis and inflammatory marker measurements, we aimed to identify associations between SARS-CoV-2 variants, microorganism abundance, and the expression of inflammatory markers.

We implemented a three-stage pipeline consisting of: (1) establishing a baseline KNN model trained with all available descriptors, (2) determining the top-ranked features through RF feature importance analysis, and (3) retraining a second set of KNN models using only the most informative features derived from RF. Specifically, two post-RF configurations were evaluated, one using the Top 20 (Sup. Figure 2-4) and another using the Top 10 ranked features, both optimized under a LOOCV scheme.

Firstly, we explored how this pipeline could be applied to train a model capable of classifying samples between positive and negative subjects. The trained model exhibited high accuracy in distinguishing between positive and negative samples as shown in Fig. 5. The RF model ranked TNF-α expression and *Acinetobacter sp.* abundance as the most important features among others, which were used to train the second KNN model using the top 10 ranked features (Sup. Table 2), increasing the AUC from 0.864 to 0.926. The average precision (AP) of the unrefined model was 0.852, whereas the post-RF model reached an AP of 0.910. Both curves indicate a clear improvement in predictive performance after feature selection.

**Figure 5.**
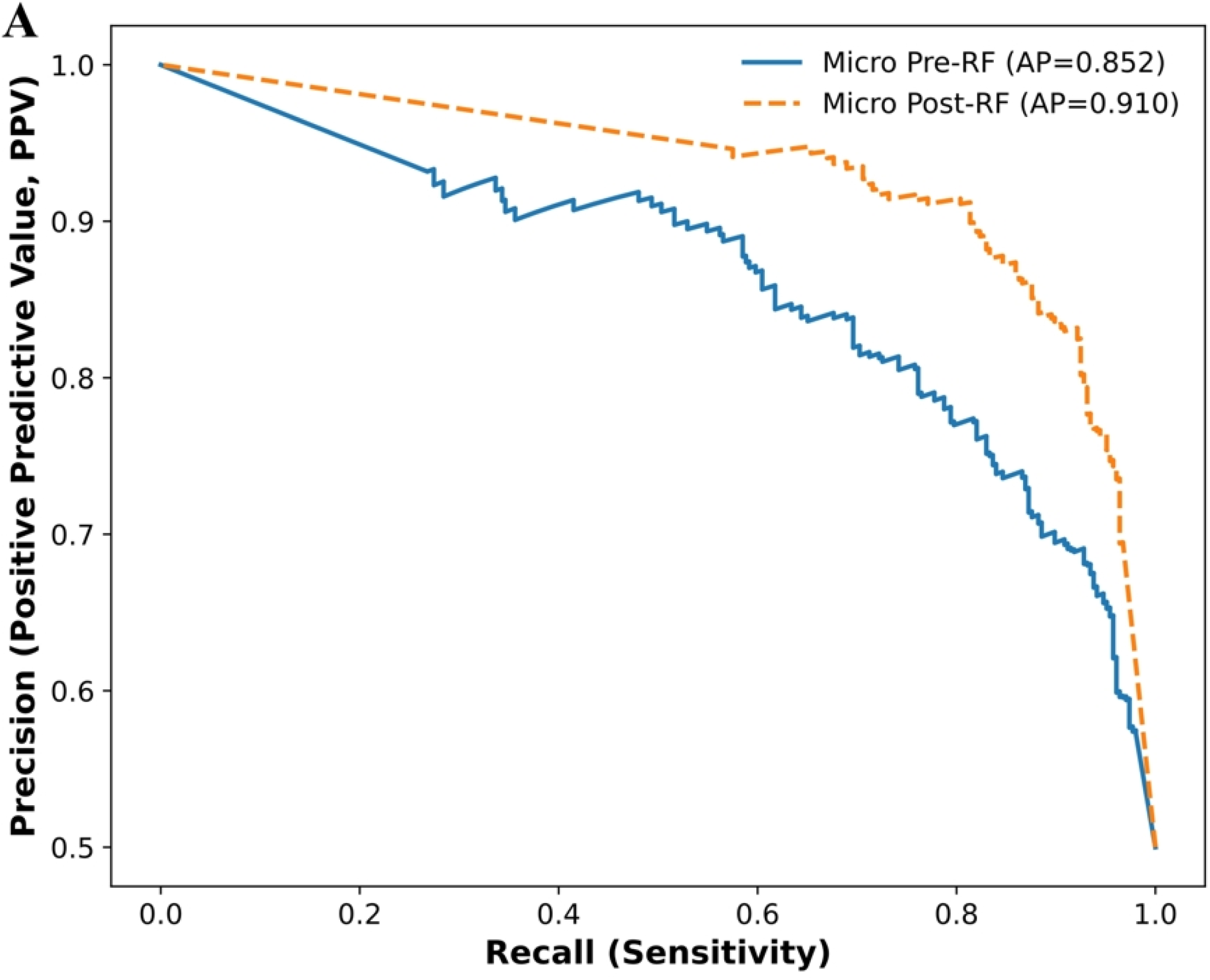
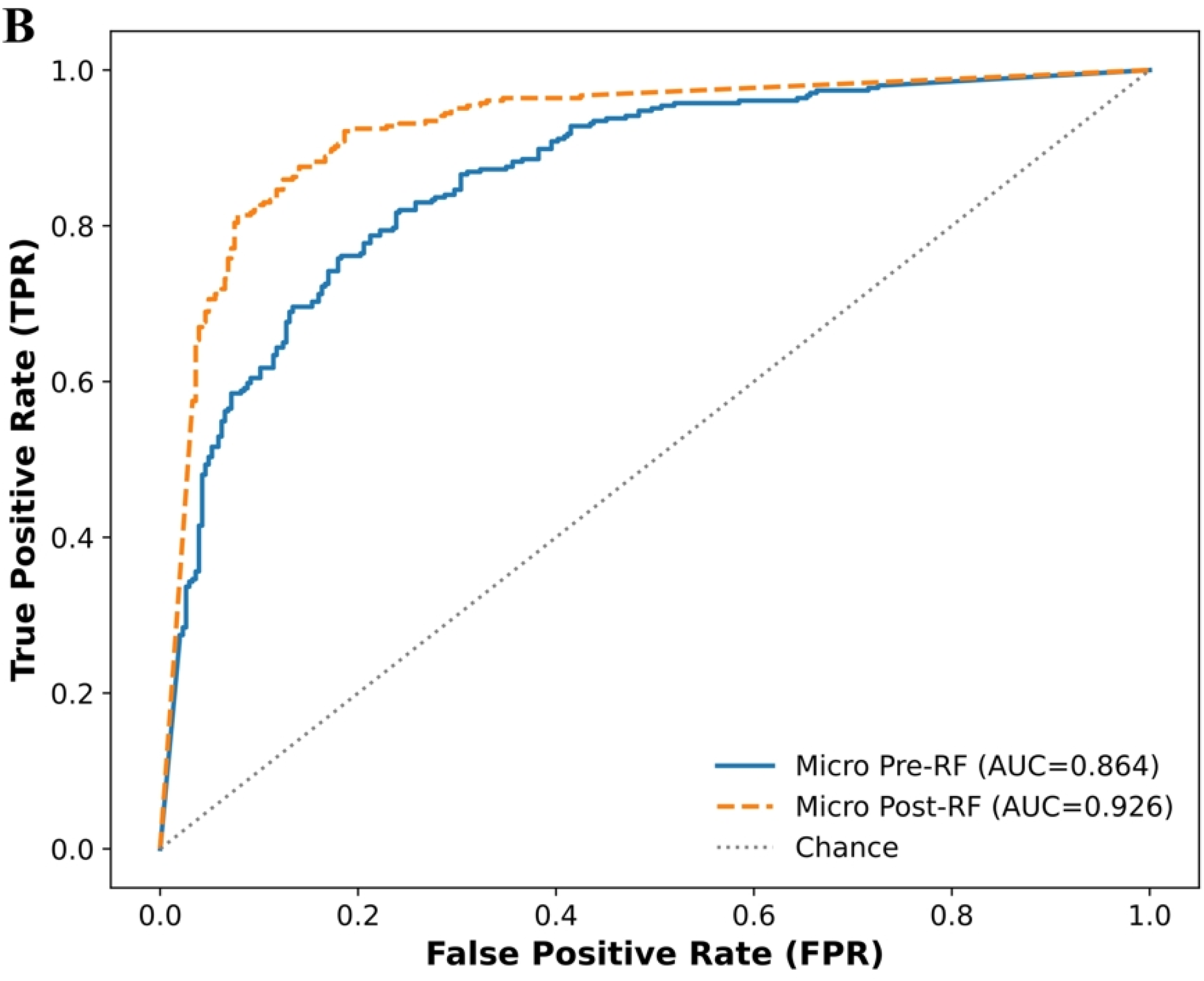
KNN model performance for SARS-CoV-2–positive and negative samples. A) PR and B) ROC weighted curves of the KNN models, showing AUC values for discrimination between positive and negative samples. Models were trained using both all available descriptors and the top-ranked features identified by RF.

Furthermore, the model was extended to classify all samples as either an individual SARS-CoV-2 variant or as negative controls. In this multiclass configuration, the baseline KNN trained with all descriptors achieved an Area Under the Curve (AUC) of 0.697 in the Receiver Operating Characteristic (ROC) curve and an averaged precision (AP) of 0.460, indicating limited discrimination and substantial overlap among Delta, Gamma, Lambda, and negative groups. When retraining the model using only the Top 10 features ranked by the RF model (Sup. Table 3), overall performance showed a slight decline, with ROC-AUC decreasing to 0.690 and AP to 0.425 (Fig. 6).

**Figure 6.**
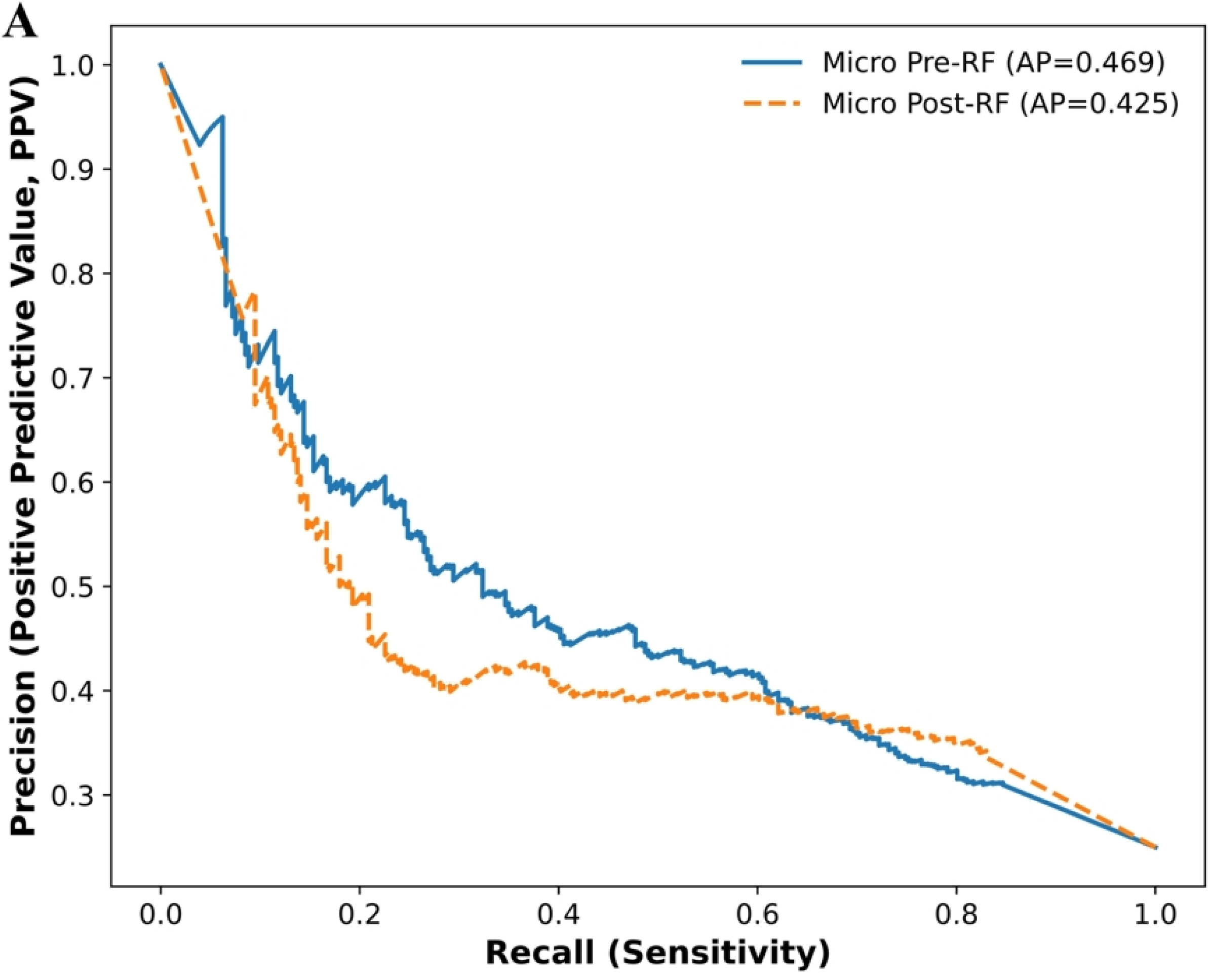
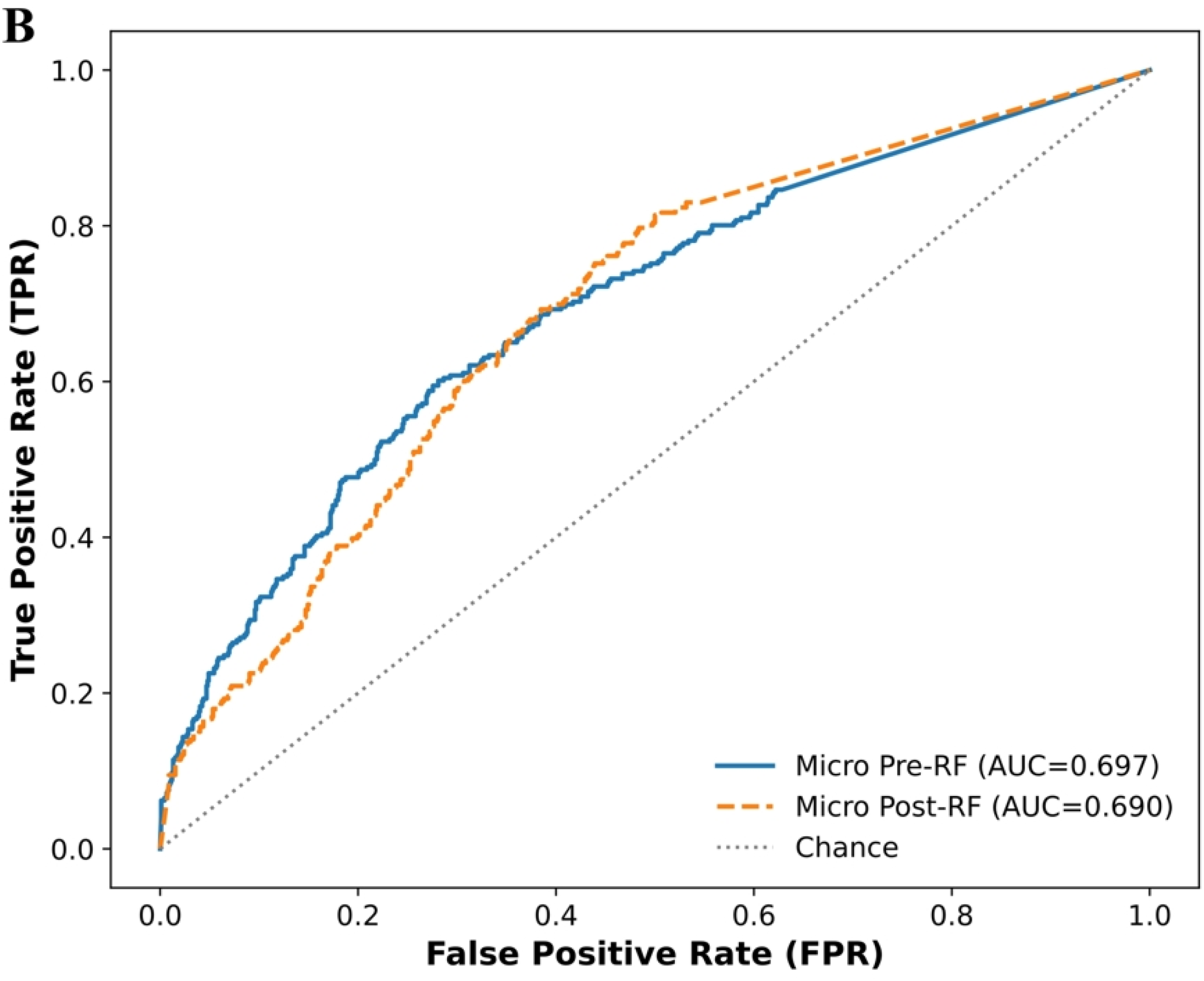
KNN model performance for SARS-CoV-2 variants and negative samples. A) PR and B) ROC weighted curves of the KNN models showing AUC values for the discrimination between SARS-CoV-2 variants (Gamma, Lambda and Delta) and negative samples. Models were trained using both all available descriptors and the top-ranked features identified by RF.

We next explored how the model would perform when only two of the three viral variants (Lambda and Gamma) were considered for classification against negative samples. The resulting model exhibited a modest improvement in overall discriminatory performance. A benchmark KNN model trained with all descriptors achieved an average ROC-AUC of 0.725 and an AP of 0.594. Applying RF feature selection to identify the most informative taxa and retraining the model with the top 10 features (Sup. Table 4) resulted in essentially equivalent performance (ROC-AUC of 0.743, AP of 0.572). This suggests that while COVID-19-associated dysbiosis is undeniably multifactorial, a fundamental core subset of features can account for most of the discriminative signal observed. These results are illustrated in Fig. 7, which displays the precision–recall and ROC curves for models trained with all descriptors and with the top-ranked features.

**Figure 7.**
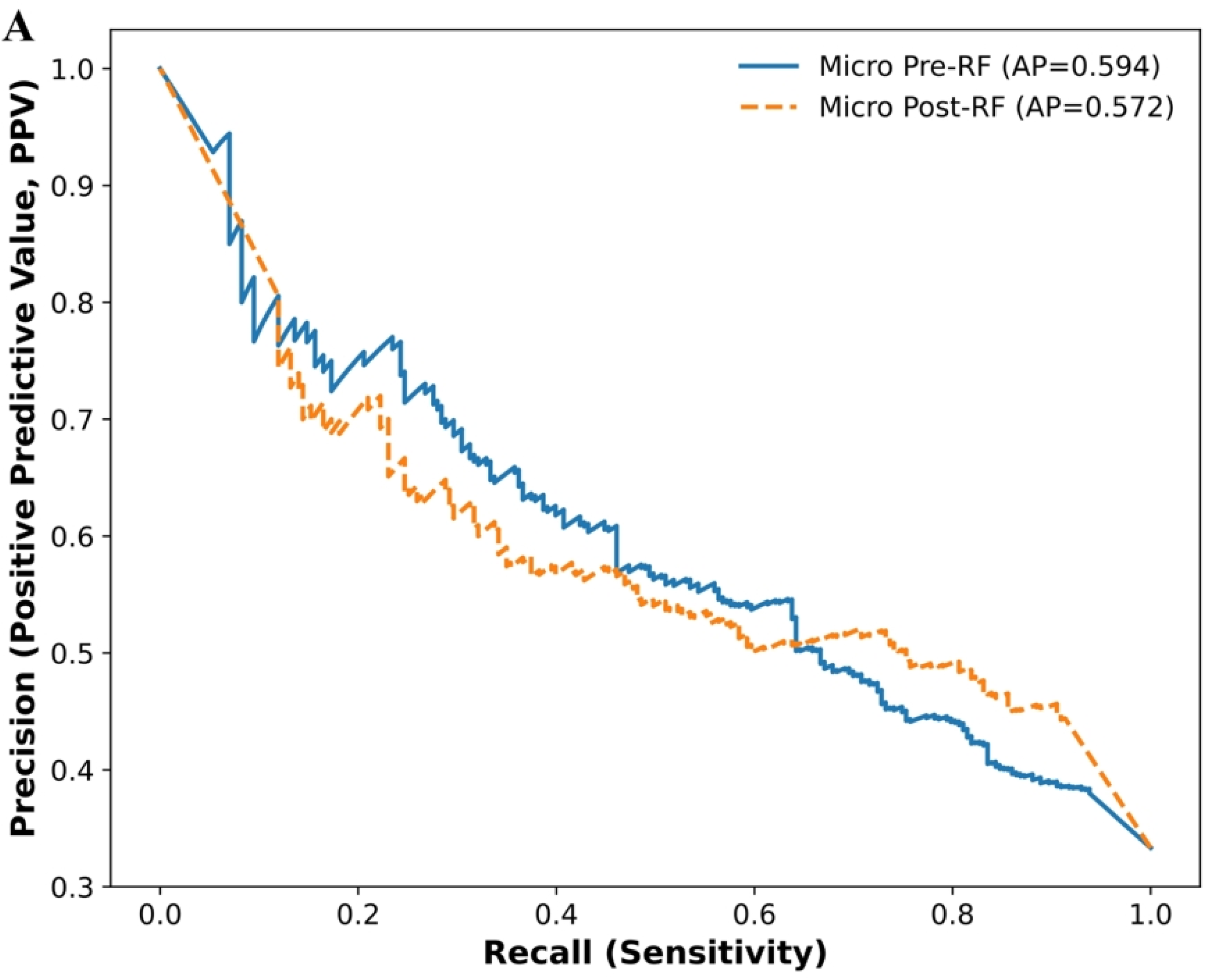
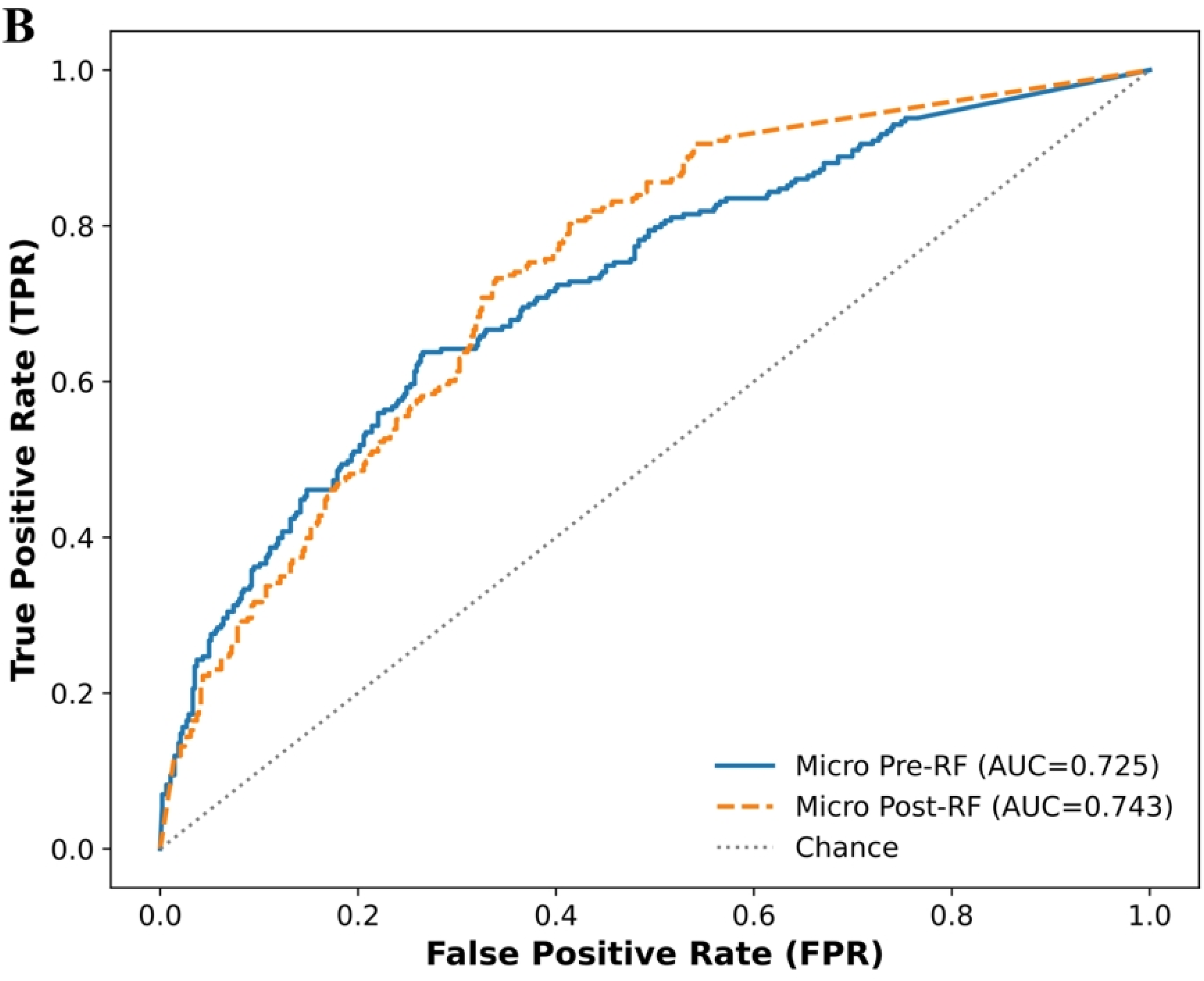
KNN model performance for SARS-CoV-2 variants Gamma, Lambda and negative samples. A) PR and B) ROC weighted curves of the KNN models, showing AUC values for the discrimination between positive and negative samples. Models were trained using both all available descriptors and the top-ranked features identified by RF.

## Discussion

Our results show that SARS-CoV-2 infection disrupts the organization of the respiratory microbiome, regardless of the viral variant involved. While microbial richness remains similar between infected and healthy individuals (Chao1 index), the Shannon and Simpson indices show that infected patients have a more even microbial community. This unexpected increase in diversity likely reflects the loss of protective dominance by key commensal bacteria, which in healthy airways help maintain ecological balance. When this structure collapses during infection, it creates space for opportunistic species to expand^9,45^. The same pattern across all variants suggests that respiratory dysbiosis is a fundamental feature of SARS-CoV-2 infection rather than a variant-specific effect. The decline in Proteobacteria and the simultaneous rise in Firmicutes and Bacteroidetes may reflect inflammation and changes in the mucosal environment triggered by the virus. At the genera level, the loss of Pseudomonas dominance and the growth of opportunistic bacteria such as *Acinetobacter sp.*, *Corynebacterium sp.*, *Moraxella sp.*, and *Staphylococcus sp.* point to a shift from a stable, commensal-rich community toward one that favors potential pathogens. This comorbidity phenomenon is largely documented across airway viral infections, which is one of the key factors in worse clinical outcomes in patients ^46,47^.

Analyses of β-diversity revealed compositional separation between positive and negative samples. Lambda, Gamma, and Delta variants exhibited extensive overlap with each other, and inclusively with the negative group, though to a lesser extent. Regarding the pro-inflammatory context, TNF-α expression was elevated in positive cases compared to negative ones, while other markers showed higher inter-individual heterogeneity and no consistent differences across variants^48^. It is important to note that, due to the contingency conditions prevailing during sample collection, the negative group may have included individuals with non–SARS-CoV-2 viral infections. This factor could explain the high variance observed in inflammatory marker expression within this group^9^. Given that TNF-α showed the most significant difference between positive and negative samples, we hypothesized that its relative expression could play a key role in the subsequent training of our classification models.

These prior results prompted an exploration into the relevance of this conglomerate of variables for training various ML models. Our initial model demonstrated high accuracy (0.864) in classifying positive versus negative samples. Upon retraining with only the top-ranked features obtained from the RF model, the resulting KNN model exhibited a significant increase in accuracy. This indicates that these descriptors collectively provide the most relevant contributions, enhancing the model’s generalization capabilities. This finding aligns with previous observations reporting TNF-α as a central cytokine in SARS-CoV-2 induced inflammation^16^ and Acinetobacter as an opportunistic pathogen frequently associated with secondary infections in COVID-19 patients^49^.

When all three viral variants (Lambda, Gamma, and Delta) were included in the classification task alongside negative samples, overall accuracy declined (AUC = 0.697). The Delta variant was often misclassified, overlapping with both negatives and other variants. This pattern mirrors the biological observation that Delta samples displayed microbiota compositions with fewer genera associated with opportunistic pathogens compared with other variants, possibly due to reduced virulence and widespread vaccination during its circulation^50,51^.

To improve the discriminatory performance, we retrained the KNN model using only the top ranked RF features under the same conditions. This approach led to a moderate loss in performance (AUC = 0.690) demonstrating that in this context a reduced selection of features does not improve the performance of our KNN models. To determine the influence of the Delta variant in the reduction of the performance of our previous KNN models, we retrained our data excluding Delta samples, focusing only on Lambda and Gamma versus negatives. In this configuration, the overall predictive performance increased (AUC = 0.726). These results indicate that the Delta variant acted as a masking class in our cohort, reducing separability among groups due to its attenuated dysbiosis and inflammatory response. The consistent improvement in AUC following class reduction highlights how differences in inflammation markers and opportunistic pathogen abundance can directly influence computational model accuracy.

Across all models, the most informative descriptors—TNF-α, IL-2, *Acinetobacter*, *Staphylococcus*, and *Prevotella*—constitute a recognizable group of specific markers linked to SARS-CoV-2 infections. Elevated TNF-α expression combined with the expansion of opportunistic genera such as *Acinetobacter* sp. and *Staphylococcus* sp. supports the notion that inflammatory activation promotes opportunistic colonization and secondary infection. This is further supported by recent studies showing that SARS-CoV-2 infection predisposes patients to bacterial coinfection with Staphylococcus aureus, which can exacerbate disease severity even when the bacterial isolates display low cytotoxicity^52^. Meanwhile, variability in IL-2 expression across individuals reflects heterogeneous immune activation, consistent with reports describing the dual role of IL-2 in antiviral protection and cytokine storm-related complications^18^. The enrichment of Prevotella observed in more severe COVID-19 cases further supports a dysbiotic shift toward anaerobic and pro-inflammatory genera in the upper airway, as previously reported in nasopharyngeal microbiome studies of infected subjects^53^. Collectively, these findings reinforce that SARS-CoV-2 infection triggers a combination of microbial imbalance and host inflammatory signaling, where the interaction of cytokines such as TNF-α and IL-2 with opportunistic bacterial taxa defines the immunomicrobial profile underlying infection severity and symptomatology across variants.

## Conclusions

The results presented in this study have demonstrated that relative abundance of opportunistic pathogens and inflammatory markers allow discrimination between SARS-CoV-2 positive and negative patients, and between individual SARS-CoV-2 variants to a lesser degree with TNF-α expression and *Acinetobacter* abundance identified as key features. The ML models implemented in our study are capable of effectively differentiating between the Gamma and Lambda variants, while the Delta variable is more susceptible to misclassification.

## Data Availability Statement

The sequencing data generated in this study will be deposited in the NCBI Gene Expression Omnibus (GEO) under accession number [to be assigned]. Additional data and analysis scripts are available from the corresponding authors upon reasonable request.

## Plain Language Summary

COVID-19 is caused by the SARS-CoV-2 virus, which mainly infects the respiratory tract. In this study, we investigated how different SARS-CoV-2 variants affected both the bacteria in the nasopharyngeal tract and the immune response of infected subjects in a Chilean cohort. We combined bacterial DNA sequencing and cytokines expression with machine learning models. We found that the virus changes the balance of bacteria in the nasopharyngeal tract, promoting the growth of certain opportunistic genera such as *Acinetobacter sp.*, *Prevotella sp.*, and *Staphylococcus sp.*. These changes were linked to higher levels of inflammation, especially of TNF-α. Our findings show how the virus, bacteria, and immune system interact during infection, and how machine learning can help identify biological patterns that may guide future treatments for respiratory diseases.

## Acknowledgements

The authors extend their gratitude to the Metropolitan Health Services (Occidente, Sur-Oriente, Sur, and Norte), the Metropolitan Regional Ministerial Secretariat of Health (SEREMI), DIGERA (División de Gestión de Redes Asistenciales), and the Ministry of Health of Chile. This acknowledgment is for the opportunity to participate jointly in the detection and diagnosis of SARS-CoV-2 in the Metropolitan Region, a collaboration that was fundamental to the development of this study. Powered@NLHPC: This research was partially supported by the supercomputing infrastructure of the NLHPC (CCSS210001).

## Disclosure Statement

The authors declare no conflicts of interest.

## Funding

This work was supported by Universidad San Sebastián (USS) under grant [USS-FIN-25-COVI-02] (Waldo A. Díaz-Vásquez), and [FB210008], Financiamiento Basal para Centros Científicos y Tecnológicos de Excelencia, Agencia Nacional de Investigación y Desarrollo (Centro Ciencia & Vida - Alberto J.M. Martin).

## References

1. Khare S, GISAID core curation team, et al. GISAID’s Role in Pandemic Response. China CDC Wkly. 2021; 3(49): 1049–1051. doi: 10.46234/ccdcw2021.255

2. Chaudhary N, Weissman D, Whitehead KA. mRNA vaccines for infectious diseases: principles, delivery and clinical translation. Nat Rev Drug Discov. 2021;20:817–38. doi:10.1038/s41573-021-00283-5.

3. Bourgonje AR, Abdulle AE, Timens W, et al. Angiotensin-converting enzyme 2 (ACE2), SARS-CoV-2 and the pathophysiology of coronavirus disease 2019. J Pathol. 2020;251(3):228–48. doi:10.1002/path.5471.

4. Yang J, Zheng Y, Gou X, Pu K, Chen Z, Guo Q, et al. Prevalence of comorbidities and its effects in patients infected with SARS-CoV-2: a systematic review and meta-analysis. Int J Infect Dis. 2020;94:91–5. doi:10.1016/j.ijid.2020.03.017.

5. Bashir A, Li S, Ye Y, Zheng Q, Knanghat R, Bashir F, et al. SARS-CoV-2 S protein harbors a furin cleavage site located in a short loop between antiparallel â-strands. Int J Biol Macromol. 2024;281:136020. doi:10.1016/j.ijbiomac.2024.136020.

6. Xie Y, Du D, Karki CB, et al. Revealing the mechanism of SARS-CoV-2 spike protein binding with ACE2. Comput Sci Eng. 2020;22(6):21–9. doi:10.1109/MCSE.2020.3015511.

7. Jain J, Gaur S, Chaudhary Y, et al. The molecular biology of intracellular events during coronavirus infection cycle. Virusdisease. 2020;31(2):1–5. doi:10.1007/s13337-020-00591-1.

8. Valdebenito S, et al. Viral titers and tissue tropism of SARS-CoV-2 in the respiratory tract. Front Microbiol. 2021;12:642934. doi:10.3389/fmicb.2021.642934.

9. Claus J, Top J, Paganelli FL, et al. Nasopharyngeal microbiome composition by SARS-CoV-2 presence and severity. Sci Rep. 2025;15:23185. doi:10.1038/s41598-025-01764-y.

10. Bustos IG, Wiscovitch-Russo R, Singh H, et al. Major alteration of lung microbiome and host responses in critically ill COVID-19 patients with high viral load. Sci Rep. 2024;14:27637. doi:10.1038/s41598-024-78992-1.

11. Hoque MN, Sarkar MMH, Rahman MS, et al. SARS-CoV-2 infection reduces human nasopharyngeal commensal microbiome with inclusion of pathobionts. Sci Rep. 2021;11:24042. doi:10.1038/s41598-021-03245-4.

12. De Maio F, Posteraro B, Ponziani FR, et al. Nasopharyngeal microbiota profiling of SARS-CoV-2-infected patients. Biol Proced Online. 2020;22:18. doi:10.1186/s12575-020-00131-7.

13. Bose T, Wasimuddin Acharya V, Pinna NK, Kaur H, Ranjan M, et al. A cross-sectional study on the nasopharyngeal microbiota of individuals with SARS-CoV-2 infection across three COVID-19 waves in India. Front Microbiol. 2023;14:1238829. doi:10.3389/fmicb.2023.1238829.

14. Gu L, Deng H, Ren Z, et al. Dynamic changes in the microbiome and mucosal immune microenvironment of the lower respiratory tract by influenza virus infection. Front Microbiol. 2019;10:2491. doi:10.3389/fmicb.2019.02491.

15. Cardinale V, Capurso G, Ianiro G, et al. Intestinal permeability changes with bacterial translocation as key events modulating systemic host immune response to SARS-CoV-2. Dig Liver Dis. 2020;52(12):1383–9. doi:10.1016/j.dld.2020.09.009.

16. Karki R, Sharma BR, Tuladhar S, et al. Synergism of TNF-á and IFN-ã triggers inflammatory cell death, tissue damage, and mortality in SARS-CoV-2 infection and cytokine shock syndromes. Cell. 2021;184(1):149–168.e17. doi:10.1016/j.cell.2020.11.025.

17. Thümmler L, Gäckler A, Bormann M, et al. Cellular and humoral immunity against different SARS-CoV-2 variants is detectable but reduced in vaccinated kidney transplant patients. Vaccines (Basel). 2022;10(8):1348. doi:10.3390/vaccines10081348.

18. Ghanbari Naeini L, Abbasi L, Karimi F, et al. The important role of interleukin-2 in COVID-19. J Immunol Res. 2023;2023:7097329. doi:10.1155/2023/7097329.

19. Larsen JM. The immune response to Prevotella bacteria in chronic inflammatory disease. Immunology. 2017;151:363–74. doi:10.1111/imm.12760.

20. Xu P, Ji X, Li M, et al. Small data machine learning in materials science. npj Comput Mater. 2023;9:42. doi:10.1038/s41524-023-01000-z.

21. Witten IH, Frank E, Hall MA. Data Mining: Practical Machine Learning Tools and Techniques. 3rd ed. Burlington (MA): Morgan Kaufmann; 2011. doi:10.1016/C2009-0-19715-5

22. Liu KJ, Zelazowska MA, McBride KM. Longitudinal analysis of convergent antibody VDJ regions in SARS-CoV-2-positive patients using RNA-seq. Viruses. 2023;15(6):1253. doi:10.3390/v15061253.

23. Li P, Luo H, Ji B, et al. Machine learning for data integration in human gut microbiome. Microb Cell Fact. 2022;21:241. doi:10.1186/s12934-022-01973-4.

24. Unal M, Bostanci E, Ozkul C, et al. Crohn’s disease prediction using sequence-based machine learning analysis of human microbiome. Diagnostics 2023;13(17):2835. doi:10.3390/diagnostics13172835.

25. Liu Y, Zhang P, Sheng H, et al. 16S rRNA gene sequencing and machine learning reveal correlation between drug abuse and human host gut microbiota. Addict Biol. 2023;28(10):e13311. doi:10.1111/adb.13311.

26. Hajeebu S, Ngembus NJ, Bandi PS, et al. Machine learning as a tool in investigating the possible role of microbiome in development and treatment of cancer. Cureus. 2021;13(8):e17415. doi:10.7759/cureus.17415.

27. Walsh I, Fishman D, Garcia-Gasulla D, et al. DOME: recommendations for supervised machine learning validation in biology. Nat Methods. 2021;18:1127–34. doi:10.1038/s41592-021-01205-4.

28. Kuncheva LI, Rodríguez JJ. On feature selection protocols for very low-sample-size data. Pattern Recognit. 2018;81:660–73. doi:10.1016/j.patcog.2018.03.012.

29. Vargas AA, Cisterna BA, Saavedra-Leiva F, et al. On biophysical properties and sensitivity to gap junction blockers of connexin 39 hemichannels expressed in HeLa cells. Front Physiol. 2017;8:243. doi:10.3389/fphys.2017.00038.

30. Van’t Veer LJ, Dai H, van de Vijver MJ, et al. Gene expression profiling predicts clinical outcome of breast cancer. Nature. 2002;415(6871):530–6. doi:10.1038/415530a.

31. Wang R, Dai W, Gong J, et al. Development of a combined nomogram integrating deep learning-pathomics, radiomics and immunoscore to predict postoperative outcome of colorectal-cancer lung-metastasis patients. J Hematol Oncol. 2022;15(1):11. doi:10.1186/s13045-022-01225-3.

32. Koh DM, Papanikolaou N, Bick U, et al. Artificial intelligence and machine learning in cancer imaging. Commun Med. 2022;2:133. doi:10.1038/s43856-022-00199-0.

33. Odom A, Varki R, Johnson W. MetaScope: Tools and functions for preprocessing 16S and metagenomic sequencing microbiome data, version 1.1.4; 2023. Available from: https://github.com/odomlab/MetaScope.

34. Parada AE, Needham DM, Fuhrman JA. Every base matters: assessing small subunit rRNA primers for marine microbiomes with DADA2. Environ Microbiol. 2016;18(5):1403–14. doi:10.1111/1462-2920.13023.

35. Love MI, Huber W, Anders S. Moderated estimation of fold change and dispersion for RNA-seq data with DESeq2. Genome Biol. 2014;15:550. doi:10.1186/s13059-014-0550-8.

36. Quast C, Pruesse E, Yilmaz P, et al. The SILVA ribosomal RNA gene database project: improved data processing and web-based tools. Nucleic Acids Res. 2013;41:D590–6. doi:10.1093/nar/gks1219.

37. Lahti L, Shetty S. microbiome: microbiome analysis package for R. Version 1.9.21. 2012– 2019. Available from: https://microbiome.github.io

38. Breiman L. Random forests. Mach Learn. 2001;45(1):5–32. doi:10.1023/A:1010933404324.

39. Bertolini R, Finch SJ, Nehm RH. Enhancing data pipelines for forecasting student performance: integrating feature selection with cross-validation. Int J Educ Technol High Educ. 2021;18:44. doi:10.1186/s41239-021-00279-6.

40. Pedregosa F, Varoquaux G, Gramfort A, et al. Scikit-learn: machine learning in Python. J Mach Learn Res. 2011;12:2825–30. Available from: https://www.jmlr.org/papers/v12/pedregosa11a.html

41. Chen K, Weng R, Li J, et al. Dual threat: susceptibility mechanisms and treatment strategies for COVID-19 and bacterial co-infections. Comput Struct Biotechnol J. 2025;27:2107–22. doi:10.1016/j.csbj.2025.05.033.

42. Gupta A, Karyakarte R, Joshi S, et al. Nasopharyngeal microbiome reveals the prevalence of opportunistic pathogens in SARS-CoV-2-infected individuals and their association with host types. Microbes Infect. 2022;24:104880. doi:10.1016/j.micinf.2021.104880.

43. Ferrari L, Favero C, Solazzo G, et al. Nasopharyngeal bacterial microbiota composition and SARS-CoV-2 IgG antibody maintenance in asymptomatic/paucisymptomatic subjects. Front Cell Infect Microbiol. 2022;12:882302. doi:10.3389/fcimb.2022.882302.

44. Smail SW, Babaei E, Amin K, et al. Serum IL-23, IL-10, and TNF-á predict in-hospital mortality in COVID-19 patients. Front Immunol. 2023;14:1145840. doi:10.3389/fimmu.2023.1145840.

45. Edouard S, Million M, Bachar D, et al. The nasopharyngeal microbiota in patients with viral respiratory tract infections is enriched in bacterial pathogens. Eur J Clin Microbiol Infect Dis. 2018;37(9):1725–1733. doi:10.1007/s10096-018-3305-8

46. Kardaras FS, Siatravani E, Tsilipounidaki K, et al. Identification of nasopharyngeal microbial dysbiosis in COVID-19 patients by 16S rRNA gene sequencing. Front Microbiol. 2025;16:1631198. doi:10.3389/fmicb.2025.1631198.

47. Marrella V, Nicchiotti F, Cassani B. Microbiota and Immunity during Respiratory Infections: Lung and Gut Affair. Int J Mol Sci. 2024;25(7):4051. doi:10.3390/ijms25074051

48. Merchant M, Ashraf J, Masood KI, et al. SARS-CoV-2 variants induce increased inflammatory gene expression but reduced interferon responses and heme synthesis compared with wild-type strains. Sci Rep. 2024;14:76401. doi:10.1038/s41598-024-76401-1.

49. Pérez-Sanz F, Tyrkalska SD, Álvarez-Santacruz C, et al. Age- and disease-severity-associated changes in the nasopharyngeal microbiota of COVID-19 patients. iScience. 2025;28:112091. doi:10.1016/j.isci.2025.112091.

50. Zhu X, Gebo KA, Abraham AG, et al. Dynamics of inflammatory responses after SARS-CoV-2 infection by vaccination status in the USA: a prospective cohort study. Lancet Microbe. 2023;4(9):e692–e703. doi:10.1016/S2666-5247(23)00171-4.

51. Wroñski J, Massalska M, Jaszczyk B, et al. Beyond interferon gamma – decreased cellular response to COVID-19 vaccination booster in patients with autoimmune inflammatory rheumatic diseases. Front Immunol. 2025;16:1568439. doi:10.3389/fimmu.2025.1568439.

52. Lubkin A, Bernard-Raichon L, DuMont AL, et al. SARS-CoV-2 infection predisposes patients to coinfection with Staphylococcus aureus. mBio. 2024;15(4):e01667–24. doi:10.1128/mbio.01667-24.

53. Ventero MP, Cuadrat RRC, Vidal I, et al. Nasopharyngeal microbial communities of patients infected with SARS-CoV-2 that developed COVID-19. Front Microbiol. 2021;12:637430. doi:10.3389/fmicb.2021.637430.

